# Jointly analyzing the association of human milk nutrients with cognition and temperament traits during the first 6 months of life

**DOI:** 10.1101/2022.04.24.489325

**Authors:** Tengfei Li, Tinu M. Samuel, Ziliang Zhu, Brittany Howell, Seoyoon Cho, Kristine Baluyot, Heather Hazlett, Jed T. Elison, Di Wu, Jonas Hauser, Norbert Sprenger, Hongtu Zhu, Weili Lin

## Abstract

Early dietary exposure via human milk (HM) components offers a window of opportunity to support cognitive and temperamental development. While several studies have focused on associations of few pre-selected HM components with cognition and temperament, it is highly plausible that HM components synergistically and jointly support cognitive and behavioral development in early life. We aimed to discern the combined associations of a wide array of HM nutrients with cognition and temperament during the first six months of life and explore if there were persistent effects up to 18 months old, when HM is the primary source of an infant’s nutrition. The Mullen Scales of Early Learning and Infant Behavior Questionnaires-Revised were used to assess cognition and temperament, respectively, of fifty-four exclusively/predominantly breastfed infants in the first 6 months of life, whose follow-ups were conducted at 6-9, 9-12 and 12-18 months old. HM samples were obtained from the mothers of the participants at less than 6 months of life and analyzed for fatty acids (total monounsaturated fatty acids, polyunsaturated fatty acid, total saturated fatty acid (TSFA), arachidonic acid (ARA), docosahexaenoic acid (DHA), ARA/DHA, omega-6/omega-3 polyunsaturated fatty acids ratio (n-6/n-3)), phospholipids (phosphatidylcholine, phosphatidylethanolamine (PE), phosphatidylinositol (PI), sphingomyelin) and choline (free choline, phosphocholine (PCho), glycerophosphocholine). Feature selection was performed to select nutrients associated with cognition and temperament, respectively. The combined effects of selected nutrients were analyzed using multiple regression. A positive association between the arachidonic acid (ARA) and surgency was observed (*p* = 0.024). Significant effect of DHA, n-6/n-3, PE and TSFA concentrations on receptive language (*R*^*2*^ = 0.39, *p* = 0.025), and the elevated ARA, PCho, and PI with increased surgency (*R*^*2*^ = 0.43, *p* = 0.003) was identified, suggesting that DHA and ARA may have distinct roles for temperament and language functions. Furthermore, the exploratory association analyses suggest that the effects of HM nutrients on R.L. and surgency may persist beyond the first 6 months of life, particularly surgency at 12-18 months (*p* = 0.002). Our studies highlighted that various HM nutrients work together to support the development of cognition and temperament traits during early infancy.

## 1 INTRODUCTION

Human milk (HM) has been considered the best and primary source of nutrients for infants particularly during the first 6 months of life. HM contains a wide variety of nutrients, bioactive components, immunological compounds, and commensal bacteria which are vital for infants’ survival, health, immunity, brain maturation, and cognitive and temperament development (1, 2). Associations between breastfeeding (BF) and improved child cognitive functions have been widely documented (3); exclusive BF promotes infant receptive language (4) and executive function (5) later in life. BF also has a significant effect on temperament development. Children who were exclusively breastfed had higher surgency and regulation, a trait reflecting inclination towards high levels of positive affect (6), and lower negative affectivity than those who were formula-fed (7). BF duration might be negatively associated with infant fussiness and positively associated with infant unpredictability (8) although these influences on the infant may be mediated by maternal sensitivity (9).

Preclinical and human studies thus far have largely focused on three main classes of nutrients in HM, namely the fatty acids, phospholipids, and choline/choline metabolites, separately regarding their effects on early cognitive development. Docosahexaenoic acid (DHA) and arachidonic acid (ARA), two of the long-chain polyunsaturated fatty acids (LCPUFAs), are believed to be involved in infant brain structural and cognitive development (10). It was reported that DHA and ARA supplementation can improve memory and problem-solving scores of preterm infants (11). In contrast, the potential effects of total saturated fatty acid (TSFA) are controversial. Some argued that increased saturated fat intake was associated with impaired ability to maintain multiple task sets in working memory and to flexibly modulate cognitive operations, particularly when faced with greater cognitive challenges (12). Phospholipids have also long been shown positively associated with brain cognitive processes (13). As major components of biological membranes and particularly abundant in the nervous system (14), phospholipids could act on the hypothalamic-pituitary-adrenal axis (13) and the microbiota-gut-brain axis (15) to exert their beneficial effects on the brain. Choline provides substrates for phosphatidylcholine and sphingomyelin formation, which are essential for neuronal and other cellular membranes, potentially improving signal transduction and brain development (16). Regarding how aforementioned HM nutrients may be associated with temperament traits, few significant findings have been reported. For example, the total Omega-3 LCPUFA was reported to be associated with infant negative affectivity, but this was not significant at an individual level for DHA, Eicosapentaenoic acid (EPA), or Eicosatetraenoic acid (ETA), the three components of the Omega-3 LCPUFA (17). The endocannabinoids, a class of ARA derivatives, were shown critically important for motivational processes, emotion, stress responses, pain, and energy balance (18).

Despite the well-recognized importance of HM on infant health, most of the above studies, focused only on the effects of an individual HM nutrient or class of nutrients. Nutrients in HM form a biological system (19); single nutrient supplementations are usually overly simplistic, ignoring the existence of other nutrients and their interplays within such a system. While jointly analyzing the micronutrient effects such as vitamins and iron was performed decades ago (20), this has rarely been investigated for the HM nutrients described above. To this end, we aimed to first jointly examine the overall effects of the three major classes of HM nutrients (fatty acids, phospholipids, and choline) on cognition and temperament traits during the first 6 months of life and second to determine if the effects persist up to 18 months of age.

## 2 MATERIALS AND METHODS

A subset of infants (n=54) enrolled in the Baby Connectome Project - Enriched (BCP-E) study (21) who were exclusively/predominantly breastfed (fed less than 4 teaspoons or 20g per day of non-formula and complementary foods/liquids (water, apple juice, etc.)) and younger than 6 months old (chronological mean age: 4.43 ± 0.83 months) were included. Children were returned for follow-up visits up to 18 months of age.

Subject recruitment and data collection were conducted by two institutions (University of North Carolina at Chapel Hill, Chapel Hill, NC and University of Minnesota, Twin Cities, MN). All study activities were approved by the Institutional Review Boards of the two universities. Informed consent was obtained from parents prior to enrolling in the study. The inclusion criteria included: birth at gestational age 37–42 weeks; birth weight appropriate for gestational age; and absence of major pregnancy and delivery complications. The exclusion criteria were: adopted infant; birth weight < 2,000 grams; abnormal MR in previous MR imaging; contraindication for MRI; neonatal hypoxia (10 minute APGAR < 5); chromosomal or major congenital abnormality; illness requiring NICU stay > 2 days; significant medical illness or developmental delay, or significant medical and/or genetic conditions affecting growth, development, or cognition (including visual/hearing impairment); the presence of a first degree relative with autism, intellectual disability, schizophrenia, or bipolar disorder; or maternal preeclampsia, placental abruption, HIV status, and alcohol or illicit drug use during pregnancy.

We used R version 3.6.3 for all the following statistical analyses. We fixed significance level α = 0.05 and corrected for regression model significance with Benjamini-Hochberg False Discovery Rate (FDR) adjustment.

### 2.1 Human Milk Collection and Macronutrient Analyses

HM samples were collected at each visit from the second feed of the day whenever possible and from the right breast using a hospital-grade, electric Medela Symphony breast pump. Mothers were instructed to completely express the contents of their right breast to ensure that the collected HM was representative of nutrients received by the infant across a full feed. The collected HM samples were vortexed at maximum speed for 2 minutes, whose volume and weight were measured and recorded with special attention to avoid bubbles. Subsequently, 3mL of HM was used for mid-infrared spectroscopic analyses using the MIRIS Human Milk Analyzer. This step was to ensure that the total fat content (as an indicator of the quality of milk sampling) fell within the expected range. Finally, an aliquot of the minimum 30mL of volume was transferred from the collection bottle to a 50mL glass beaker. Using a repeating pipette and an appropriate tip, eleven aliquots of 1mL were made in 1mL Eppendorf tubes, and nine aliquots of 2mL were made in 2mL Eppendorf tubes for storage in a −80 °C freezer within 30 minutes from the end of the time of collection.

### 2.2 Fatty acids, Phospholipids and Choline Analyses

A representative 1mL aliquot of collected HM was shipped on dry ice to Nestle Research Center (Switzerland) for analyses of fatty acids (FAs) and phospholipids whereas choline was analyzed at UNC-Chapel Hill. Specifically, direct quantifications of FAs were accomplished using gas chromatography as detailed in (22) and the total monounsaturated fatty acids (TMUFA), total polyunsaturated fatty acids (TPUFA), TSFA, ARA, DHA, and ARA to DHA ratio (ARA/DHA), Omega-6/omega-3 polyunsaturated fatty acids ratio (n-6/n-3) were obtained. In contrast, analyses of phospholipids were accomplished using high-performance liquid chromatography coupled with a mass spectrometer detector (23), which yielded phosphatidylcholine (PC), phosphatidylethanolamine (PE), phosphatidylinositol (PI), and sphingomyelin (SPH). Finally, quantification of choline and choline metabolites was performed using liquid chromatography-stable isotope dilution-multiple reaction monitoring mass spectrometry (LC-SID-MRM/MS). Chromatographic separations were performed on an Acquity HILIC 1.6 μm 2.1×50mm column (Waters Corp, Milford, USA) using a Waters ACQUITY UPLC system and free choline, phosphocholine (PCho), and glycerophosphocholine (GPC) were obtained.

### 2.3 Assessments of Cognition and Temperament

Two measures, namely the Mullen Scales of Early Learning (MSEL) and Infant Behavior Questionnaires-Revised (IBQ-R) (23), were employed to assess cognition and temperament, respectively. The MSEL includes five subscales: fine motor (F.M.), gross motor (G.M.), visual reception (V.R.), receptive language (R.L.), and expressive language (E.L.). An early learning composite (E.L.C) score considered as the Developmental Quotient of infants was calculated by summing the T-scores of all subdomains excluding G.M. The MSEL was administered by trained staff at every visit. In contrast, the IBQ-R is a widely used 14-scale parent report measure designed to assess infant temperament. To reduce the number of variables, three factors were extracted by using three linear weighted averages of the 14 subscales. Specifically, we incorporated the weights of three latent factors reported in the exploratory factor analysis results of Gartstein and Rothbart (24), and obtained three personal traits, namely, surgency/extraversion (SUR), negative affectivity (NEG), and orienting/regulation (REG). The subscale items included in the above three factors and loading scales are provided in **Supplementary Table 1**. In our study, the MSEL or IBQ-R assessments in the first 6 months were obtained within 30 days from the collection of HM samples.

### 2.4 Statistical Modeling for Associations Analyses in the First 6 Months of Life

All HM nutrient concentrations were first normalized with mean zero and unit variance. Marginal association analysis between each individual nutrient and each subdomain of MSEL and IBQ-R scores were carried out by a linear regression of each subdomain score on each nutrient. Since nutrients could vary with postpartum duration, to ensure that age does not contribute to our analyses, age was included as a controlled variable. Other confounding factors including sex, data collection site and household income (if < 75k) were also controlled in the above linear model and the standardized regression coefficient was obtained between each nutrient and each MSEL/IBQ-R score. Household incomes of two infants were missing and imputed with the overall average.

To evaluate the potential combined and conditional effects of HM nutrients in association with MSEL and IBQ-R, and to deal with the collinearity among the HM nutrients, the correlation matrix among all nutrients was calculated. Subsequently, nutrients were clustered into several subgroups based on their correlation using the single linkage clustering analysis (SLCA) (25). The SLCA approach was chosen since it offers intuitive interpretations of the results; all pairs of nutrients exhibiting correlation coefficients greater than a predefined threshold T_C_ were combined into one sub-group. The optimal number of clusters was determined by maximizing the Dunn index (26). Furthermore, in order to minimize redundancy and collinearity (27), the best subset selection model (28) was employed by selecting at most one nutrient from each cluster with the highest adjusted R^2^. The chosen nutrients were then used as the main effects and MSEL or IBQ-R obtained within 30 days before or after the collection of HM samples as the outcomes for the regression model to uncover the potential associations. The overall significance was reported using the ANOVA F statistics in comparison to the reduced baseline model. More detailed description on SLCA and the statistical models are relegated to **Supplementary information 4** and **5**. To assess if the identified nutrients predict cognitive and temperament scores of new subjects, multiple regression models with the above selected nutrients were evaluated by squared cross validation (CV) errors and prediction correlations through 100 repetitions of five-fold CV.

### 2.5 Model evaluation on follow-up visits beyond 6 months

To evaluate if the identified associations of HM components and cognition/temperament persisted beyond the first 6 months of life in the same subjects, subjects with follow-up visits were binned into three age groups: 6-9 months, 9-12 months and 12-18 months. Each observation within each age group corresponded to a unique subject. The regression models trained above (as in **Supplementary Table 5**) were used to predict MSEL and IBQ-R scores (predicted scores), respectively. The correlation and p-value based on Pearson’s t-test between the predicted and observed scores were evaluated.

## 3 RESULTS

Of the 54 subjects, cognition was assessed using MSEL in 38 subjects (4.64 ± 0.89 months; 12 males) whereas temperament using IBQ-R in 42 subjects (4.48 ± 0.73 months; 16 males) and both were available in 26 subjects (4.81 ± 0.71 months; 9 males). After the initial assessments, follow-up assessments of MSEL were available in 27, 34, and 25 subjects at 6-9 (7.63 ± 0.87 months; 9 males), 9-12 (10.5 ± 0.97 months; 10 males) and 12-18 months old (14.02 ± 1.08 months; 8 males), respectively. In contrast, IBQ-R was obtained from 7, 7, and 27 subjects between 6∼9 (mean: 243.9±12.3 days), 9∼12 (mean: 341.1±18.8 days) and 12∼18 (mean: 13.34 ± 0.79 months) months, respectively (**Figure 1**; **Table 1**). Detailed demographic information on infant and their mothers including infant age, sex, anthropometrics, measured breast milk gross composition like fat, carbohydrates, proteins and energy, as well as household income and mother’s education are summarized in **Table 1**, respectively.

**Figure 1.**
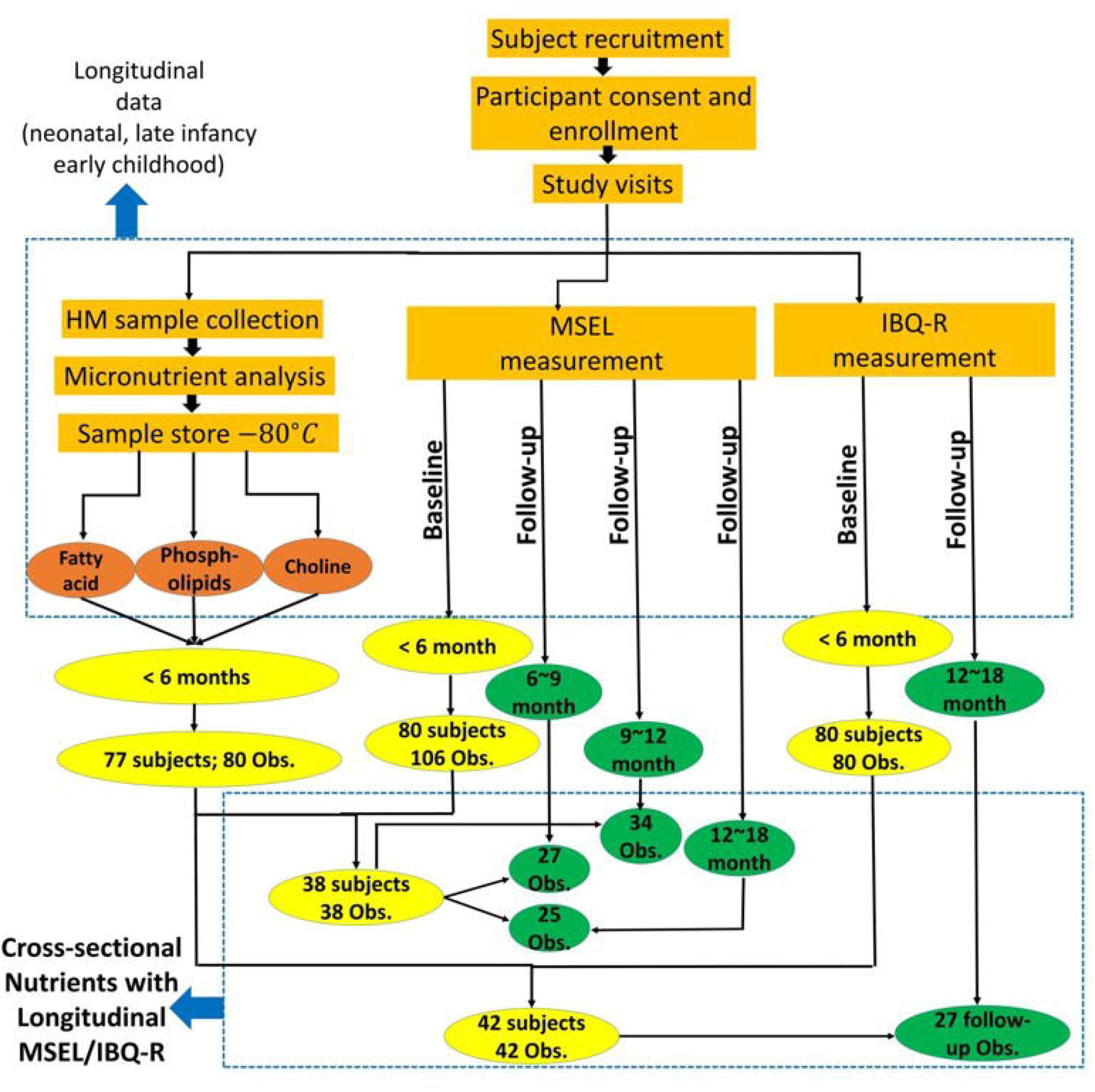
Study flowchart provides the overall experimental details.

**Table 1:**
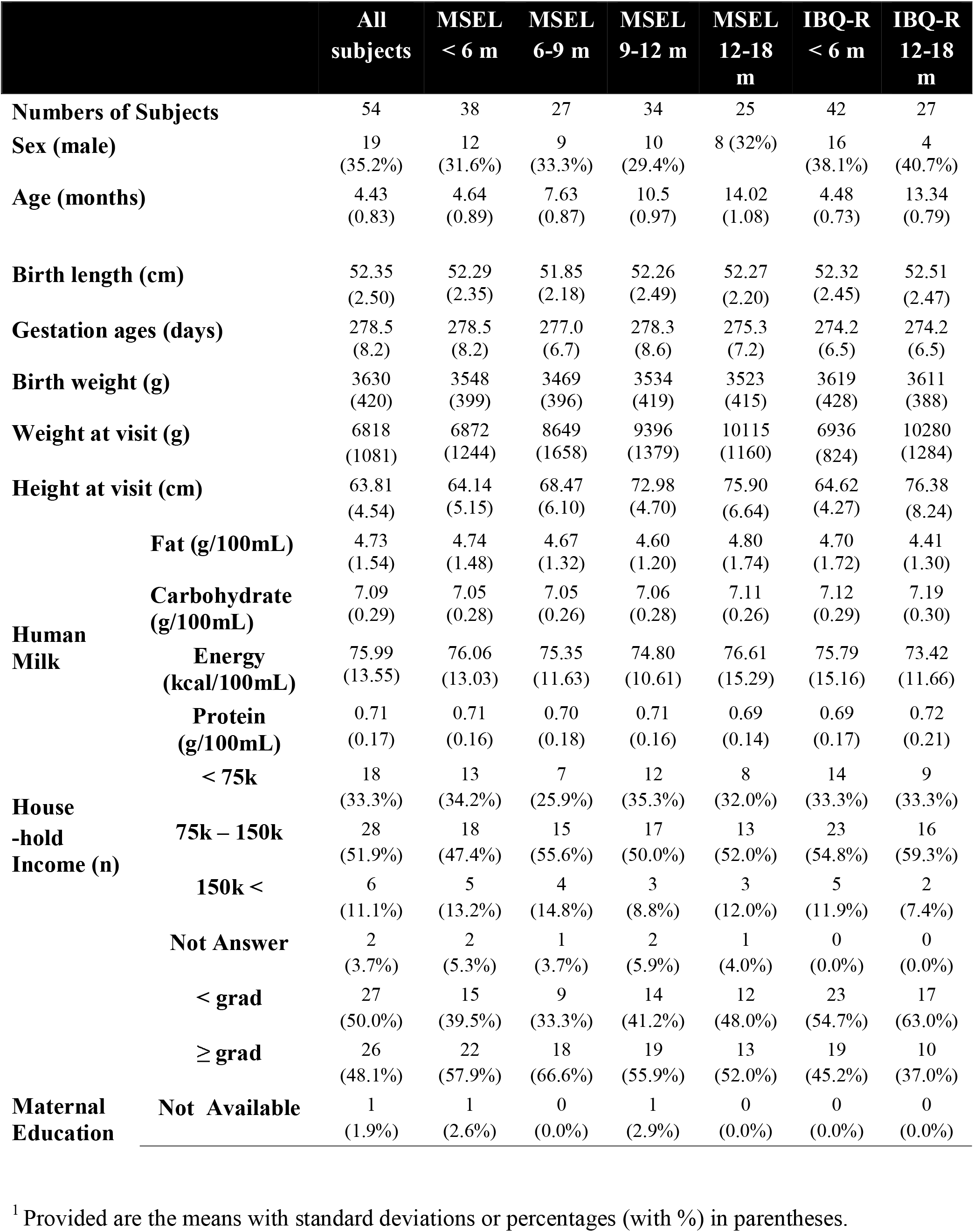
Characteristics of the participants and HM samples^**1**^

The scatterplot of the 14 nutrients is shown in **Figure 2** and their individual concentrations are summarized in **Table 2**. Among these nutrients, DHA decreases (*p* < 0.002) while ARA/DHA ratio (p < 0.03) increases with postpartum age. Since some of the subjects only had MSEL but not IBQ-R or vice versa during the first 6 months of life, the mean values of each nutrient from the MSEL dataset and the IBQ-R dataset were compared using the two-sided two-sample t-test. No significant differences were observed (**Supplementary Figure 1**; all raw *p* ≥ 0.4), suggesting that there is no selection bias of the mean nutrient concentrations. In addition, the MSEL and IBQ-R scores are provided in **Supplementary Tables 2 and 3**, respectively, which are within the normal ranges of both scales.

**Figure 2.**
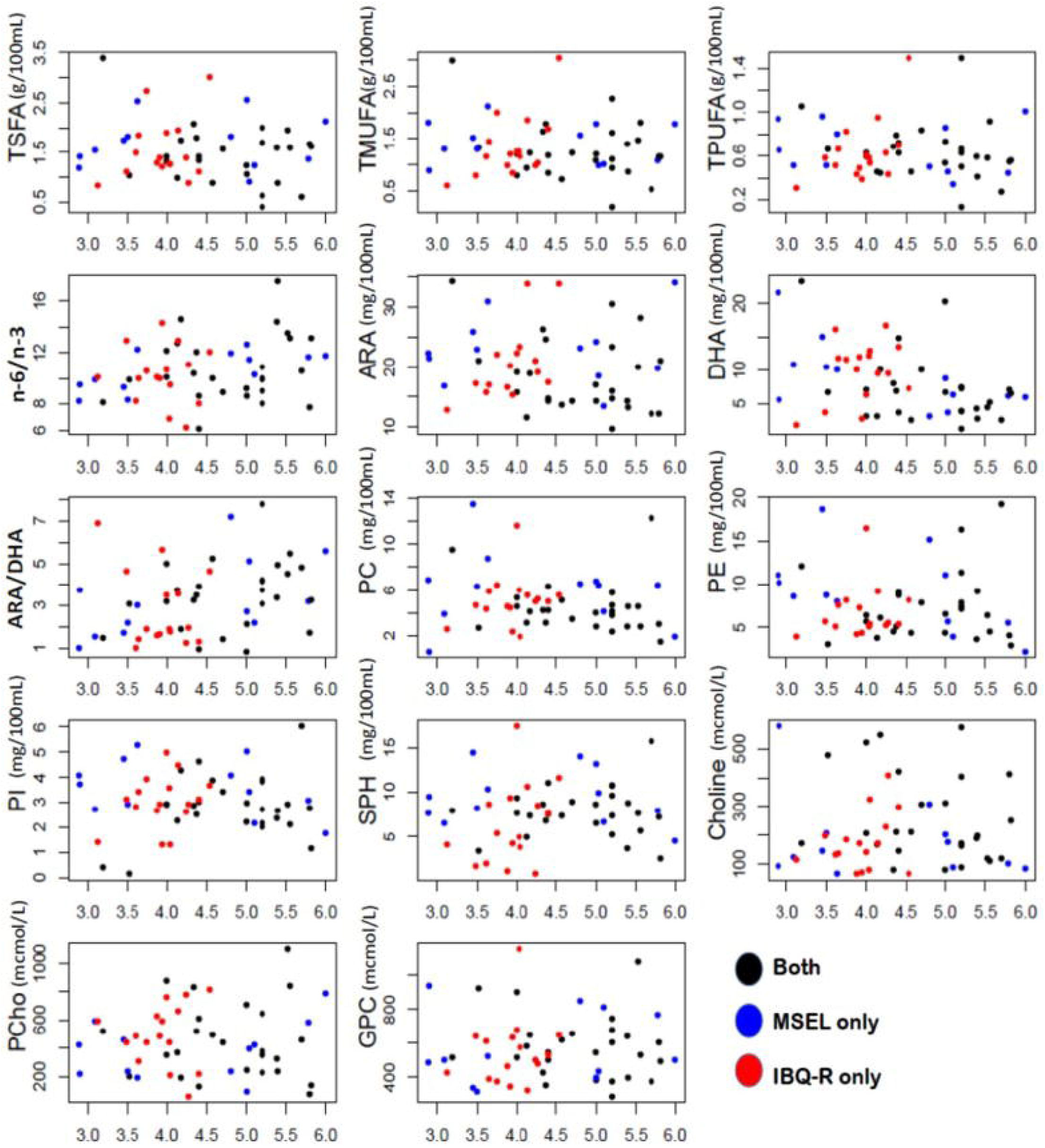
Scatterplots of the 14 nutrients with age in months (x-axis) for subjects assessed with MSEL (blue circles), IBQ-R (red circles) and both (black circles), respectively. TMUFA, total monounsaturated fatty acids; TPUFA, polyunsaturated fatty acid; TSFA, saturated fatty acid; ARA, arachidonic acid; DHA, docosahexaenoic acid; ARA/DHA, the ARA-to-DHA ratio; n-6/n-3, the Omega-6/Omega-3 polyunsaturated fatty acids ratio; PC, phospholipids; PE, phosphatidylethanolamine; PI, phosphatidylinositol; SPH, sphingomyelin; Choline, free choline; PCho, phosphocholine; GPC, Glycerophosphocholine; SUR, surgency; NEG, negative affectivity; REG, regulation.

**Table 2:** Concentration values of the 14 HM nutrients in the first 6 postpartum months

| Nutrients | Mean | Median | Std. | (Lower) 5% | (Upper) 95% |
| --- | --- | --- | --- | --- | --- |
| TSFA (g/100ml) | 1.54 | 1.43 | 0.59 | 0.76 | 2.64 |
| TMUFA (g/100ml) | 1.32 | 1.21 | 0.53 | 0.69 | 2.19 |
| TPUFA (g/100ml) | 0.64 | 0.6 | 0.26 | 0.33 | 1.03 |
| n-6/n-3 | 10.57 | 10.14 | 2.27 | 7.39 | 14.37 |
| ARA (mg/100ml) | 20.07 | 19.15 | 6.32 | 12.21 | 33.99 |
| DHA (mg/100ml)* | 8.35 | 7.09 | 5.14 | 2.58 | 18.14 |
| ARA/Total Fat (%) | 0.60 | 0.60 | 0.15 | 0.40 | 0.80 |
| DHA/Total Fat (%) | 0.24 | 0.19 | 0.14 | 0.11 | 0.52 |
| ARA/DHA* | 3.24 | 3.2 | 1.73 | 1.01 | 6.15 |
| PC (mg/100ml) | 4.99 | 4.6 | 2.58 | 1.93 | 10.51 |
| PE (mg/100ml) | 7.53 | 6.4 | 3.98 | 3.45 | 16.39 |
| PI (mg/100ml) | 3.06 | 2.94 | 1.19 | 1.24 | 5 |
| SPH (mg/100ml) | 7.7 | 7.66 | 3.61 | 1.86 | 14.19 |
| Choline (mcmol/l) | 216.49 | 173.5 | 140.47 | 72.5 | 530.45 |
| PCho (mcmol/l) | 448.85 | 439.56 | 235.64 | 120.85 | 834.8 |
| GPC (mcmol/l) | 568.53 | 528.62 | 192.07 | 335.08 | 931.9 |

\*: indicates significant changes with postpartum age

While the main focus of our study was to jointly analyze a wide array of HM nutrients in association with cognition and temperament in typically developing children, the marginal associations of each nutrient with MSEL and IBQ-R scores are shown in **Table 3** (the heatmap is shown in **Supplementary Figure 2**). A significant association between ARA and SUR (*r* = 0.54, *p* = 0.0006, adjusted *p* = 0.024, 95% CI = [0.25, 0.83]) was observed. Several other nutrients including TSFA (*r* = 0.35, *p* = 0.035, 95% CI = [0.03, 0.68]), TPUFA (*r* = 0.35, *p* = 0.031, 95% CI = [0.03, 0.67]) and PCho (*r* = 0.46, *p* = 0.003, 95% CI = [0.17, 0.76]) was positively associated with SUR; TPUFA (*r* = 0.32, *p* = 0.046, 95% CI = [0.007, 0.64]), and ARA (*r* = 0.39, *p* = 0.015, 95% CI = [0.08, 0.70]) was associated with REG. In addition, marginal negative associations between G.M. and TSFA (standardized regression coefficient *r* = -0.40, *p* = 0.01 and 95% confidence interval (CI) = [-0.71, - 0.08]) and TMUFA (*r* = -0.38, *p* = 0.02 and 95% CI = [-0.70, -0.05]) and a positive association between DHA and R.L. (*r* = 0.38, *p* = 0.049 and 95% CI = [0.002, 0.76]) were observed, although all of these associations did not pass the FDR control due to limited sample size.

**Table 3:**
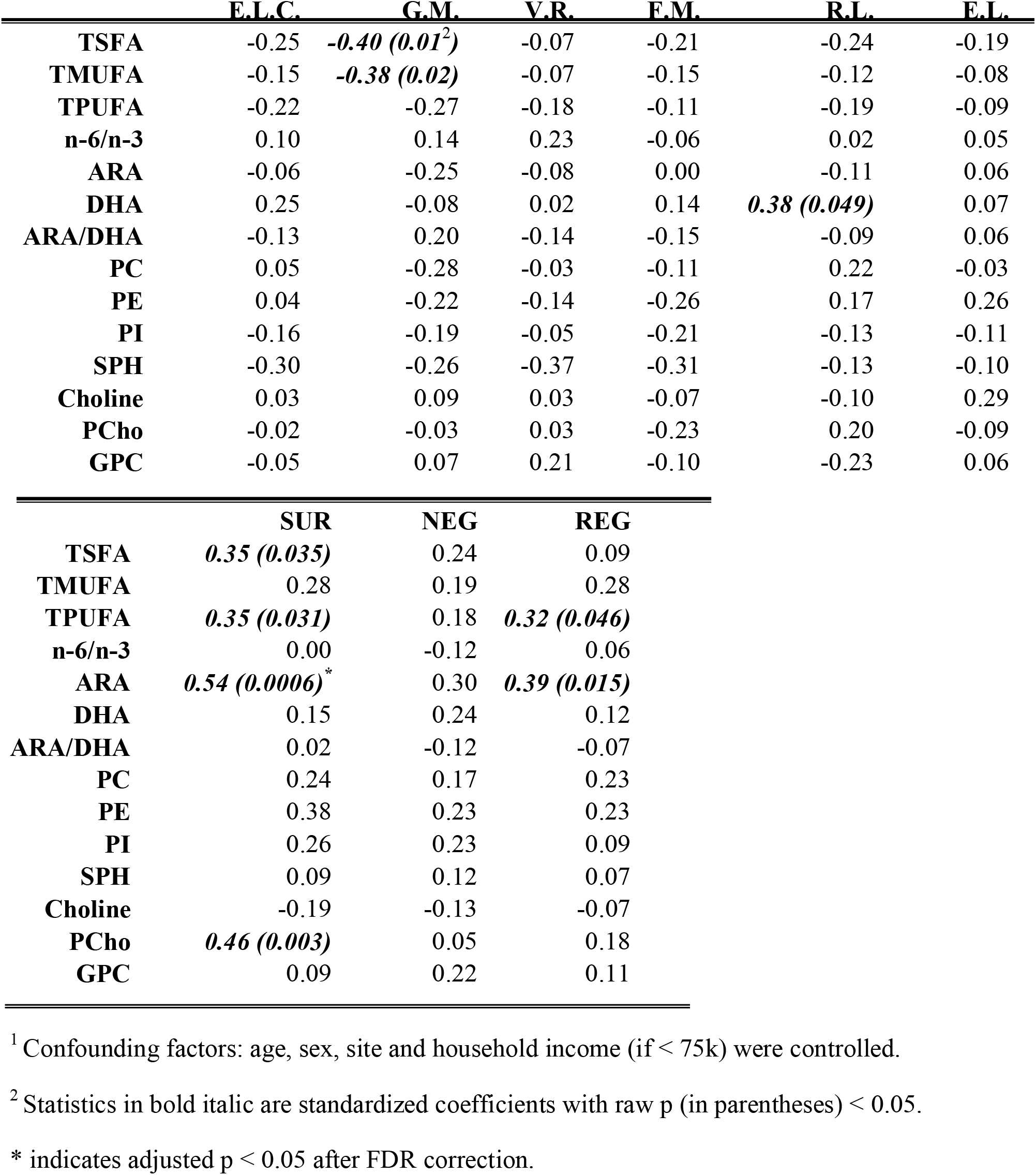
Standardized regression coefficients of the HM nutrients with the MSEL (top) and IBQ-R factors (bottom), respectively^1^.

We further tested if HM nutrients are correlated. Evidently, many of the HM nutrients were highly correlated (**Figure 3A** and **Supplementary Table 4**). In particular, the nutrients in the phospholipid family were highly correlated. As outlined above, to minimize collinearity, the single linkage clustering analysis was used to cluster highly correlated nutrients into one group and the Dunn index was used to determine the optimal number of clusters. The optimal number of cluster was 7, which exhibited the largest Dunn index (**Table 4)**. Note that the optimal clustering result was stable for 0.53 < T_C_ < 0.69. In addition, the memberships of groups 1 – 3 were stable independent of the *Tc* for 0.53 < T_C_ < 0.77. The minimum spanning tree, representing the smallest sum of distances to touch all vertices of the graph of the optimal clustering results are shown in **Figure 3B** and detailed information is given in **Supplementary information 4**.

**Figure 3.**
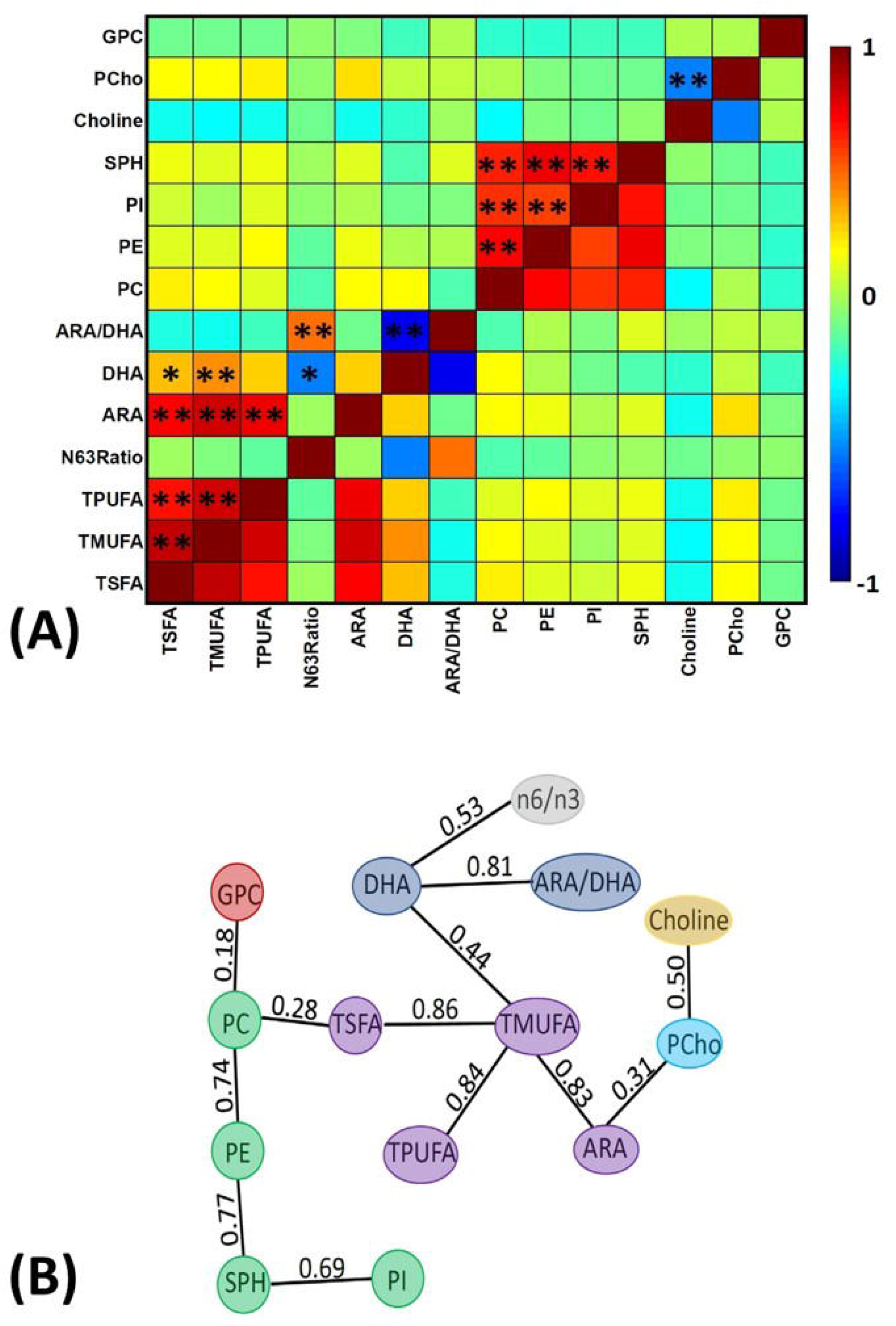
Relations between nutrients. **(A)** Pearson correlation between each pair of the 14 nutrients, where the asterisk (*) and double asterisks (**) represented a raw p < 0.05 and the adjusted p < 0.05, respectively, using the standard Pearson Correlation test and the FDR correction. **(B)** The minimum spanning tree, where colors represent different clusters and the numbers represent the correlation coefficients between a given pair of nutrient. Note the optimal clustering result is stable for 0.53 <Tc ≤ 0.69. In addition, the minimum spanning tree shows that the strongest correlation between the clusters (PC, PI, PE, SPH) and (TSFA, TMUPA, TPUFA, ARA) is between TSFA and PC (0.28). It implies that only when T_C_ < 0.28 these two clusters will merge together, while PCho and Choline are more close in the sense that these two will merge when T_C_ < 0.50.

**Table 4:** Correlation and grouping of the 14 nutrients.

|  | **Scheme 1** | **Scheme 2** | **Scheme 3** |
| --- | --- | --- | --- |
|  | *$0.53 < Tc^1 \leq 0.69$* | *$0.69 < Tc \leq 0.74$* | *$0.74 < Tc \leq 0.77$* |
| **Group 1** | TSFA, TMUFA, TPUFA, ARA | TSFA, TMUFA, TPUFA, ARA | TSFA, TMUFA, TPUFA, ARA |
| **Group 2** | n-6/n-3 | n-6/n-3 | n-6/n-3 |
| **Group 3** | DHA, ARA/DHA | DHA, ARA/DHA | DHA, ARA/DHA |
| **Group 4** | PC, PE, PI, SPH | PC, PE, SPH | PE, SPH |
| **Group 5** | GPC | Choline | Choline |
| **Group 6** | PCho | PCho | PCho |
| **Group 7** | Choline | GPC | GPC |
| **Group 8** | - | PI | PC |
| **Group 9** | - | - | PI |
| **Dunn index** | 0.64 | 0.44 | 0.41 |

<sup>1</sup>Tc: The threshold values for the correlation coefficients.

Significant combined associations of HM nutrients were observed for R.L. (MSEL) and SUR (IBQ-R) summarized in **Figure 4A**. Specifically, for receptive language, the final linear regression model includes DHA, n-6/n-3 ratio, PE and TSFA (-) and R-squared = 0.39 (adjusted *p* = 0.025), where the information provided in the parentheses indicates the sign of the coefficients and positive otherwise. In contrast, for SUR, the model includes ARA, PI and PCho with R-squared = 0.43, and adjusted *p* = 0.003. Finally, a positive association between ARA and REG with R-squared = 0.23 and adjusted *p* = 0.03 was observed. The scatter plots of the combined associations between the experimentally obtained and fitted R.L. and SUR using the identified association model are shown in **Figures 4B and 4C**, respectively. More detailed coefficients, confidence intervals, and p-values for each model are shown in **Supplementary Table 5**. In addition, the scatter plots of the associations between each of the selected nutrients for R.L., SUR, and REG are shown in **Supplementary Figure 3**. Furthermore, 5-fold CV between the observed and predicted MSEL and IBQ-R scores were conducted. The boxplots of Pearson’s correlations between the observed and predicted MSEL and IBQ-R scores from 100 random splits of 5-fold CV are shown in **Figure 5A**. The boxplots imply that in 99% and 91% cases of the 100 random splits of CV repetitions, the correlation between the observed and predicted R.L. scores and SUR were significant, respectively, whereas in most cases the association with the observed REG was not significant. These results showed a stronger prediction power of R.L. and SUR but not REG. The observed (x-axis) versus the predicted R.L. (**Figure 5B**) and SUR (**Figure 5C**) of the 100 times’ predictions are also provided. The mean and the 95% confidence interval of prediction correlations (based on the random splits of CV) were 0.434 and [0.349, 0.501] for the R.L. and 0.409 [0.281, 0.509] for the SUR. These results showed model robustness of predictions against training data perturbations.

**Figure 4.**
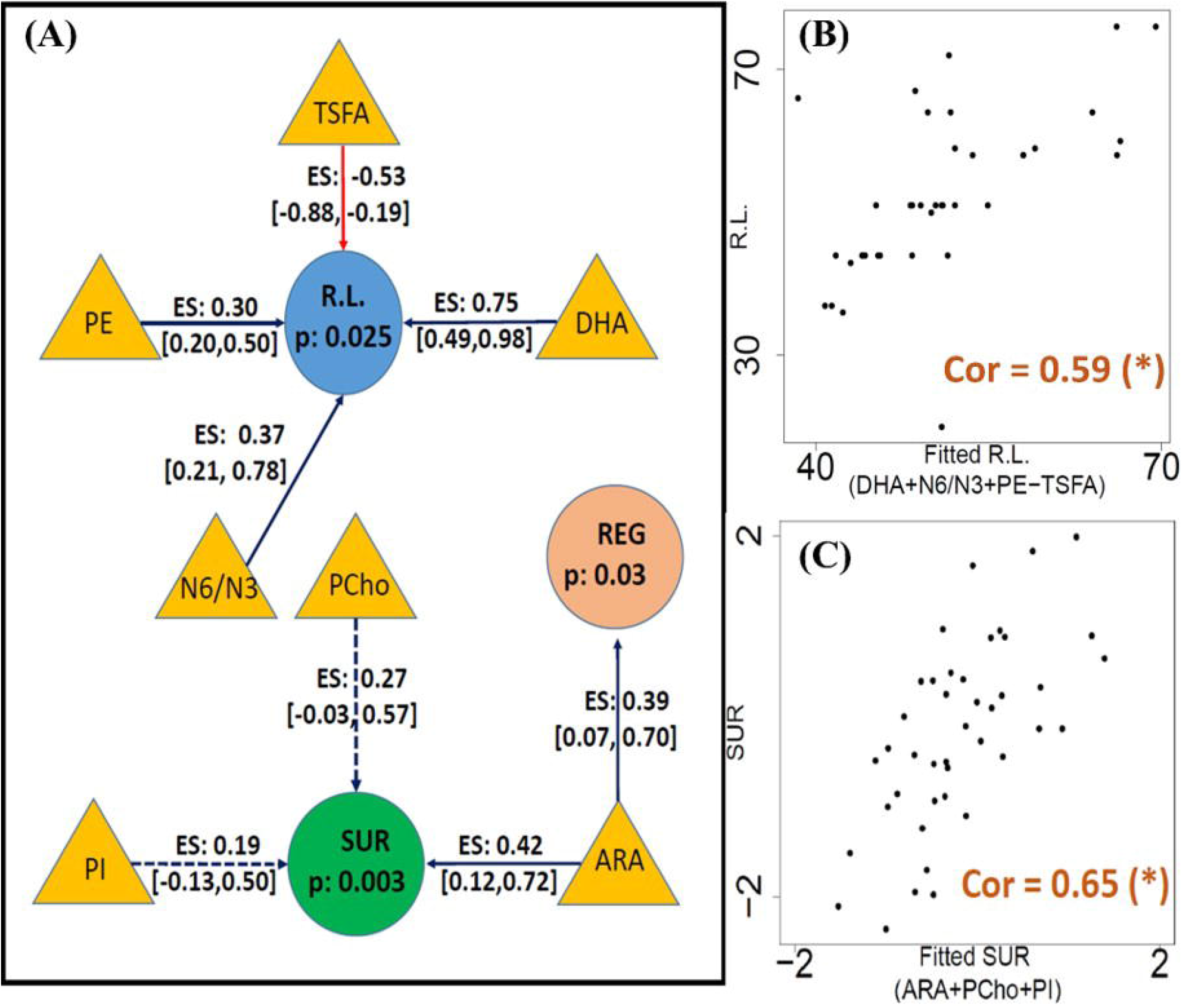
**(A)** The identified conditional of HM nutrients (filled triangles) on receptive language score (filled blue circles) and SUR (filled green circles) using linear regression. The blue and red arrows represent positive and negative associations, respectively. The corresponding effect size, confidence interval of coefficients for each association, and FDR corrected p-values for the regression model are provided. Dotted lines indicate the regression coefficient of the covariate is not significant but the inclusion of the corresponding nutrient improves model fitting with a higher adjusted R2. **(B)** The association between the experimentally obtained R.L. and the combined effect of selected HM nutrients DHA, n-6/n-3, PC, and TSFA (fitted R.L.). **(C)** The association between the experimentally obtained SUR and the combined effect of selected HM nutrients ARA, PCho, and PI (fitted SUR). Pearson’s correlation between the fitted and observed measurements are included in panels **B** and **C** with (*) indicating adjusted p < 0.05.

**Figure 5.**
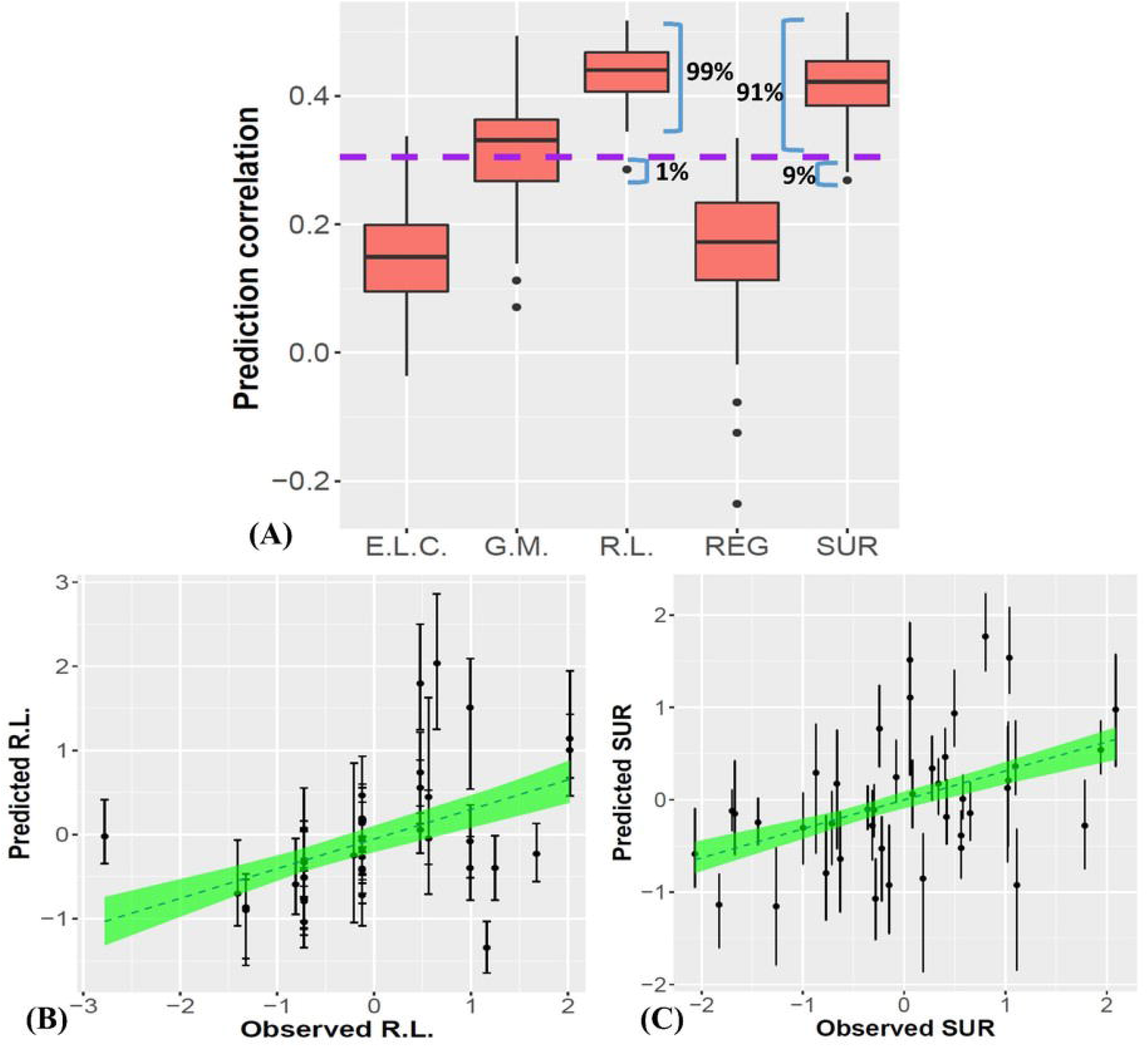
Prediction results for linear regression models from 100 random splits of 5-fold CV. **(A)** shows the boxplots of 100 Pearson’s correlations between the observed and predicted MSEL (E.L.C., G.M., and R.L.) and IBQ-R factors (REG and SUR) using linear regression, respectively, based on 100 repetitions of CVs. The purple dashed line represents the significance threshold of the correlation at α=0.05 with n=38 subjects. **(B)** and **(C)** show the observed R.L. (left) and SUR (right) versus the predicted R.L. and SUR of the 100 times’ predictions, respectively. The dashed lines and the green areas show the mean and the 95% confidence intervals of the fitted linear slopes between the observed and predicted observations.

Finally, we conducted exploratory analyses for predicting the two outcome measures at the follow-up observations using results obtained < 6 months old. With limited sample sizes for IBQ-R at the 6-9 and 9-12 months follow-up visits (**Table 1**; **Figure 1**), the IBQ-R analyses were only performed for the 12-18 months age group whereas MSEL was conducted for all the three age bins. The correlation coefficients and p-values for each age bin of MSEL and SUR at 12-18 months are provided in **Table 5**. Evidently, the associations between predicted and observed R.L. decrease with age with the best performance at 6-9 months (*p = 0*.*07*. **Figure 6A**). In contrast, a significant association between predicted and observed SUR at 12-18 months was observed (p=0.002. **Figure 6B**).

**Figure 6.**
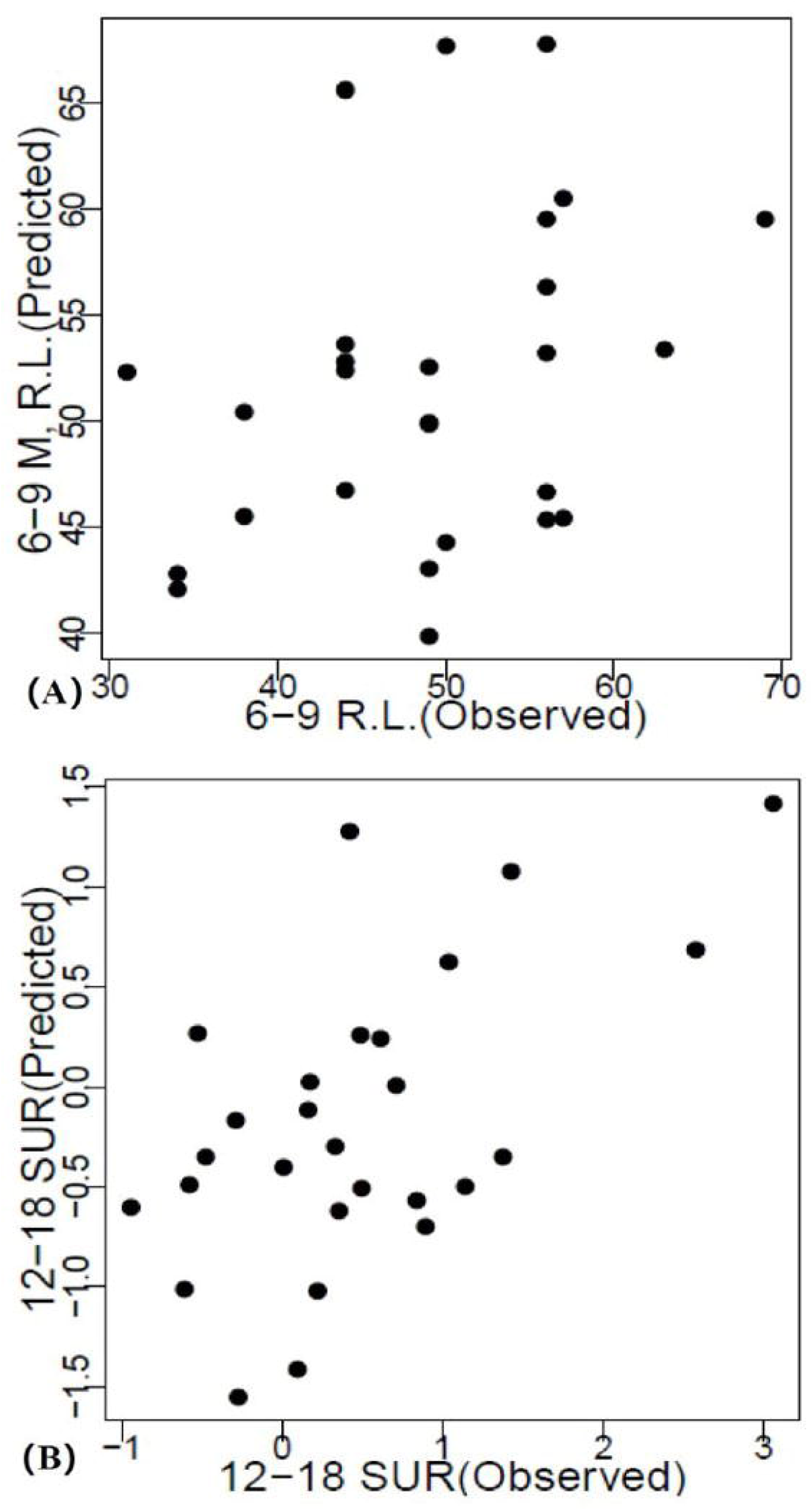
Association of the predicted and observed R.L. and SUR at the follow-up ages. **(A)** Predicted versus observed R.L. at 9-12 months. **(B)** Predicted versus observed SUR at 12-18 months.

**Table 5:** Correlation between the predicted and observed R.L. and SUR at follow-up.

| **Compare** | **RL 6-9m** |  | **RL 9-12m** |  | **RL 12-18** |  | **SUR 12-18<sup>1</sup>** |  |
| --- | --- | --- | --- | --- | --- | --- | --- | --- |
|  | *r =* | 0.35 | *r =* | 0.14 | *r =* | -0.21 | *r =* | ***0.58*** |
|  | *p =* | 0.07 | *p =* | 0.43 | *p =* | 0.32 | *p =* | ***0.002*** |

<sup>1</sup> Results with the p-values less than 0.05 are shown in italic bolded text.

## 4 DISCUSSION

The HM produced by each mother for her infant has a unique composition and contains a myriad of different lipids, vitamins, minerals, bioactive carbohydrates, proteins, and immune factors that evolve in tandem with the growth and developmental needs of the infant (29). In addition, maternal diet and lifestyle can also affect HM composition especially the lipid components (30). While several studies have shown positive associations of HM bioactives and nutrients with brain development (21, 31), most of them have largely considered one or only few nutrients. Christian, et al. (19) recently pointed out that these approaches may be overly simplistic and failed to consider the complex synergetic effects of HM nutrients on infant development and health. They advocated the need of a paradigm shift by considering HM nutrients as a biological system in future studies aiming to discern the benefits of HM nutrients. To this end, we collected HM samples, concurrently assessed cognition and temperament of BF infants, utilized advanced statistical models to jointly consider 14 widely evaluated HM nutrients spanning over the three families: choline, fatty acids, and phospholipids, and reported combined associations between HM nutrients and cognition/temperament during the first 6 months of life. Furthermore, whether the effects of identified nutrients during the first 6 months persist through to 1.5 years old were also evaluated.

### 4.1 HM nutrients in the first six postnatal months

An analysis of the temporal trends in HM nutrients over the first six postnatal months revealed that DHA exhibited a significant negative age effect and choline remained relatively stable, consistent with those reported previously (32). In contrast, while the phospholipids were stable with age, several studies have reported an increase over lactation (32, 33). HM DHA/total fat content was highly variable (ranging 0.08% to 0.71 %) with a median concentration of 0.19%, which is below the worldwide average (0.37%) and those reported from South-East Asian regions (32, 34). Nevertheless, lower levels of HM DHA/total fat have been reported in the North American region (35), possibly due to low habitual intake of seafood in the inland areas, and that levels are strongly impacted by maternal DHA intake (36). In addition, the median HM ARA/total fat of 0.60% in our cohort was higher compared to the global average (0.55 %), leading to a higher ARA/DHA ratio (3.24) when compared to that reported in the literature (1.5 – 2) (37). The HM n-6/n-3 fatty acid ratio in our study (mean 10.57), while comparable to some geographies, was higher than others (38), suggesting differences either in body fat composition and mobilization of fat stores or dietary habits such as consumption of LA-rich vegetable oils (39). Levels of most PLs (PC and PE) in our study were comparable to those reported from Singapore (32). We observed higher concentrations of free choline and GPC, but lower levels of PCho, compared to reports from Canada and Cambodia (40), possibly due to differences in dietary choline intake and amounts of choline available from the maternal circulation. Taken together and comparing with other studies, our findings highlight that several HM nutrients may show geographical differences across regions reflecting differences in diet and lifestyle.

### 4.2 Combined effects of HM nutrients on cognition and temperament

The potential interplay between nutrients and early brain development has been widely recognized (41). Our brains undergo rapid development during the first years of life by establishing new synapses (synaptogenesis), removing excessive synapses (pruning), myelination (42), and forming highly complex yet efficient brain functional networks enabling the performance of cognitive tasks and social behaviors (43). During these dynamic and highly energy-demanding brain developmental processes, appropriate nutrient supply is key for healthy neurodevelopment in infancy. Our results show that while each individual content of PE, DHA, n-6/n-3, or TSFA does not show significant association on its own (**Table 3**), a combined effect is observed on receptive language (R.L.). In contrast, ARA by itself is significantly associated with SUR (adjusted p=0.024) whereas neither PCho nor PI is (**Table 3**). Nevertheless, the three nutrients ARA, PCho, and PI combined exhibit an improved association and predictive performance with SUR in terms of elevated model significance, decreased AIC, and CV errors. Importantly, we also observed a significant association between ARA and REG. This finding differs from the marginal associations shown in **Table 3** where no associations were observed between REG and ARA after FDR correction. Largely, this finding is not surprising. Specifically, FDR correction was employed for the marginal associations to control for 14 nutrients and subscales of scores. In contrast, since multiple regression was employed with features selected using approaches outlined above, FDR correction was only employed to control for subscales of scores but not for the 14 nutrients. Instead, 5-fold cross validation was used to evaluate the possibility of overfitting. Indeed, CV results yield that the identified association between (PE, DHA, n-6/n-3, TSFA) and R.L. and (ARA, PCho, and PI) and SUR were robust but not between ARA and REG. Therefore, cautions should be taken when interpreting the association between ARA and REG. Future studies with a larger sample size are warranted to further evaluate the association between ARA and REG.

Two key findings of our study deserve additional discussion. First, while MSEL assesses five domains of cognition, significant associations were only observed between HM nutrients and receptive language in our study. Several studies have previously evaluated the potential associations between individual HM nutrients and infants’ language ability. However, thus far the results are inconsistent and sometimes controversial. Although DHA has been reported to improve language ability, a recent comprehensive review by Gawlik et al. (44) has concluded that the current evidence of DHA supplementation on language development is limited and non-conclusive. Ramos et al. (45) reported that a higher ratio of the linoleic (n6 fatty acid) to the alpha-linolenic acid (n3 fatty acid) could exert beneficial effects for R.L. in HM fed preterm infants. However, their results are contrary to the body of literature showing that n-6/n-3 ratio in the maternal blood and diet is negatively associated with vocabulary and verbal fluency of their infants (46, 47). While the effect of PE on language abilities have not been specifically reported in the literature, the Milk Fat Globule Membrane (MFGM), which is a diverse mixture of bioactive components including phospholipids, has been shown to improve visual function, language, and motor domains in both term and preterm infants (48, 49). Our results show that none of the HM nutrients when evaluated individually exhibited significant associations with infants’ language ability; however, a combined effect of PE, DHA, n-6/n-3, and TSFA is identified on R.L., underscoring the importance of considering the strong dependency between HM nutrients and the power of jointly analyzing multiple nutrients.

Second, while there is a consensus that temperament traits emerge early in life and have a strong genetic and neurobiological basis (50), what is less understood is the role of BF and HM nutrients in shaping these offspring behavioral traits. Our study highlights that HM nutrients may be associated with specific temperament traits in infancy. Notably, HM ARA (adjusted p=0.024) alone (but not DHA) and the combined effects of ARA, PCho, and PI exhibited a significant association with SUR, which includes high-intensity pleasure, smiling and laugher, perceptual sensitivity, vocal reactivity, and activity. Tallima and Ridi reported that the downstream metabolites of ARA such as eicosanoids, or endocannabinoids play a critical role in brain reward signaling, motivational processes, emotion, stress responses, and pain (18), which may further shape infants’ behavioral traits. While no study has specifically looked at HM choline levels and infant behavior, PC supplementation during pregnancy was shown to result in fewer attention problems and less social withdrawal at 40 months of infant age by normalizing the development of cerebral inhibition (51). Equally, supplementation of MFGM, rich in phospholipids, in early life was shown to be associated with fewer parent-reported behavioral problems in their children and improved behavioral regulation (52). Nevertheless, our results differ from the recent findings of Hahn-Holbrook et al (2019) who showed that higher n-3 PUFAs in HM (more specifically ALA which is a precursor of DHA), but not any of the n-6 PUFAs, was associated with significantly less sadness and distress to limitations (17). It is plausible that differences in experimental design (varying ages among subjects in our cohort vs all assessed at 3 months old by Hahn-Holbrook) and the limited sample size in our study may have contributed to the different findings. Nevertheless, comparing the HM nutrients associated with receptive language and SUR in our study, it appears that there may be distinct LCPUFA roles (DHA vs ARA) for temperament and language functions, respectively. Future studies with a larger sample size are warranted to determine if ARA or its metabolites have a distinct role from that of DHA on behavioral development.

### 4.3 Exploratory Analyses on the Effects of HM nutrients beyond the first 6 months of life

The regression models elucidating potential relations between HM components and cognition/temperament during the first 6 months of life were evaluated on subjects whose follow-up MSEL and IBQ-R were available 6-18 months of age. Using association analyses, the correlation between the predicted and observed SUR scores was significant at 12-18 months of age (*r* = 0.58, *p* = 0.002), and there is a potential association yet not significant (*p* = 0.07) between the predicted and observed R.L. at 6-9 months. These results suggests that the MSEL or SUR scores beyond 6 months can be possibly predicted using the derived regression models encompassing the combined effects of HM nutrients from the first 6 months of life. Although the limited sample sizes for SUR at the 6-9 and 9-12 months made it difficult to determine if the findings observed at 12-18 months also present during the two age periods, our results appear to suggest that the effects of HM on temperament persist longer than that of R.L., since the association of SUR is quite strong at 12-18 months, while the association of R.L. exhibits a trend toward significant during 6-9 months (p=0.07) but continues decreasing at 9-12 and 12-18 months. It is, however, worth noting that other unobserved confounders could affect the future scores, such as type of confounding factors (amount and variety), solid food intakes, and environmental stimulations. As a result, the effects of nutrients that infants received < 6 months on cognition/temperament could diminish with age. Nevertheless, our results suggest that the SUR is less affected by unobserved confounders and thus be more predictable from baseline. In-depth investigation in future studies is needed to confirm our results.

### 4.4 Conclusions

The development of cognitive and behavioral functioning is complex and likely involves the interplay of social, psychological, and biological factors. Nevertheless, HM nutrients such as LCPUFAs, choline, and phospholipids may be modifiable contributors to cognitive and behavioral development, especially during the early BF period. Our results provide evidence that specific nutrients in HM may act together to support cognitive and behavioral traits. However, these findings warrant replication in larger cohorts of BF infants with longitudinal follow-up for more definitive behavioral phenotyping, by controlling maternal diet and lifestyle, an in-depth understanding of the mechanisms by which HM nutrients jointly affect developmental trajectories, and careful examination of the synergistic effects, if any, of these nutrients on functional outcomes.

## 5 DATA AVAILABILITY STATEMENT

Data described in the manuscript, codebook, and analytic code will be made available upon request pending application and approval by the authors.

## 7 AUTHOR CONTRIBUTIONS

BH, JE and WL designed the research. KB, BH, HH, JE, and WL conducted research. TL, TS, ZZ, SC, DW and WL analyzed data or performed statistical analysis. TL, TS and WL wrote the paper. TL, TS, ZZ, SC, KB, BH, HH, JE, HZ, JH, NS, DW and WL had primary responsibility for final content. All authors have read and approved the final version of the manuscript.

## 9 ACKNOWLEDGMENTS

We acknowledge Giuffrida Francesca and Tavazzi Isabelle for analyses of HM Phospholipids. We acknowledge Neotron for the analyses of fatty acids.

