## Supplemental for "Jointly analyzing the association of human milk nutrients with cognition and temperament traits during the first 6 months of life"

Supplementary Material

### 1 Supplementary Information

#### 1.1 Supplementary Information-1: Human Milk Nutrients

The descriptive statistics of the nutrients for subjects assessed using MSEL and IBQ-R are shown in **Supplementary Figure** **1**. Using the two-sided two-sample t-test, the mean values of each nutrient from the MSEL dataset and the IBQ-R dataset have no significant difference (all raw *p* ≥ 0.4), suggesting that there is no selection bias of the mean nutrient concentrations.

#### 1.2 Supplementary Information-2: The MSEL and IBQ-R measurement

The descriptive statistics of the MSEL scores and the IBQ-R measurements (including the original 14 IBQ-R scores and the 3 IBQ-R latent factors) are provided in **Supplementary Table** **2** and **Supplementary Table** **3**, respectively. In addition, the heatmap showing the marginal associations between each nutrient and MSEL and IBQ-R is shown in **Supplementary Figure** **2.**

#### 1.3 Supplementary Information-3: Correlation and linear regression results

The correlation between each pair of the 14 nutrients is shown in **Supplementary Table** **4**, while the linear regression results for MSEL scores/IBQ-R factors with the selected nutrients are listed in **Supplementary Table** **5**. Scatterplots of the selected nutrients for receptive language score (R.L.) and surgency (SUR) are provided in (A.1)-(A.4) and (B.1)-(B.3), respectively.

#### 1.4 Supplementary Information-4: The SLCA, Minimum Spanning Tree and Dunn’s Index

The SLCA clustering, also known as the nearest neighbor clustering, is one of the most widely used approaches of hierarchical clustering. Given two sets, $X$ and $Y$, of features and a threshold $D_{C}$, the SLCA clustering can be performed by merging clusters with small single-linkage distance $D\left( X,Y \right)\leq D_{C},$where $D\left( X,Y \right)=\min_{x\in X, y\in Y} d(x,y)$ and $d\left( x,y \right)=\sqrt{2(1-cor(x,y))}$ is the correlation distance metric, $cor(x,y)$ is the Pearson correlation between features $x$and $y$, $D_{C}=\sqrt{2(1-T_{C})}$ and $T_{C}$ is a threshold of minimum correlation to determine the number of clusters. One advantage of SLCA is that it can be easily computed and understood through the minimum spanning tree. The minimum spanning tree is the spanning tree that has the smallest sum of distances to touch all vertices of the graph. As an example in Figure 3 of the main text, the strongest correlation between the clusters {PC, PI, PE, SPH} and {TSFA, TMUPA,TPUFA, ARA} is between TSFA and PC (0.28), which implies that only when $T_{C}\leq0.28$ these two clusters will merge together, while {PCho} and {Choline} are more close in the sense that these two will merge when $T_{C}\leq0.50.$

Dunn’s index returns the ratio of minimal intercluster-distance to maximal cluster diameter, denoted as$DI(\{C_{1},\ldots,C_{p}\})=\min_{1\leq i<j\leq p} D(C_{i}, C_{j})/\max_{1\leq k\leq p} \max_{x,y\in C_{k}} d(x,y)$. Dunn’s index is one of the most widely used internal indices to determine the optimal number of clusters. A higher Dunn’s index indicates that either an increased minimal intercluster-distance or a decreased maximal cluster diameter.

#### 1.5 Supplementary Information-5: Detailed statistical approaches

For marginal association analyses, we fit the following linear regression for each nutrient $X_{j},$

$Model 0: MSEL or IBQ \sim1+\sum_{k=1}^{q} C_{k}+X_{j},j=1,\ldots, p,$

where $C_{k}, k=1,\ldots, r$ are the $r$ controlled confounders.

For joint association analyses, we will apply the best subset selection method to a pool of predictor candidates (*p* = 14 nutrients). This method consists of fitting models that primarily consider every possible combination of subset covariates from the *p* predictors, with 2^p^ possible combinations in total. In our case as there is strong associations between nutrients, to reduce multi-collinearity, we only consider possible combinations from those with at most a single nutrient from each cluster. The final optimal model can be determined which maximizes the adjusted R-squared, as it adjusts the R-squared of regression to the number of terms in a model to reduce overfitting. To formularize our model, we denote $\mathcal{T}$ as the set of all the nutrients. Let $\mathcal{T}_{k}, k=1,\ldots, p$ are the $k$-th cluster obtained based on the SLCA clustering approach and $p$ is the number of clusters. Then we have $\bigcup_{k\leq p} \mathcal{T}_{k}\mathcal{=T}$ and $\mathcal{T}_{j}\cap\mathcal{T}_{k}=\emptyset$ for $j\neq k.$

$$Model 1: MSEL or IBQ \sim1+\sum_{k=1}^{r} C_{k}+\sum_{j=1}^{q} X_{j}, X_{j}\in\mathcal{T}_{k_{j}}$$

where $q$ is the number of selected nutrients which satisfies$0\leq q\leq p$, and $X_{j}$ is the selected nutrient from the $k_{j}$-th cluster $\mathcal{T}_{k_{j}}$ with $k_{j}\in\left\{ 1,\ldots, p \right\}$ and $k_{j}\neq k_{l}$ if $k\neq l$. The optimal choice of ${\{X}_{j}|j=1,\ldots p\}$ was determined by maximizing the adjusted R-squared of Model 1. We included the age, sex, data collection site and household income as confounders. The overall significance was reported using the ANOVA F statistics in comparison to the reduced model below.

$Model 2: MSEL or IBQ \sim1+\sum_{k=1}^{q} C_{k}$.

The FDR adjustment was adopted for controlling Type I errors of multiple comparisons.

### 2 Supplementary Tables and Figures

#### 2.1 Supplementary Tables

**Supplementary Table 1: IBQ-R three factor loadings (weights) and subscales included in each factor.**

| IBQ-R |  |  |  |
| --- | --- | --- | --- |
| Composite Scales | **Subdomain** | | **Loadings/weights** |
| *Negative affectivity* | high intensity pleasure | | -0.25 |
|  | sadness |  | 0.79 |
|  | distress to limitations | | 0.69 |
|  | fear |  | 0.31 |
|  | falling reactivity | | -0.56 |
| *Surgency/Extraversion* | approach |  | 0.74 |
|  | vocal reactivity | | 0.74 |
|  | high intensity pleasure | | 0.69 |
|  | smiling and laughter | | 0.55 |
|  | activity level | | 0.49 |
|  | perceptual sensitivity | | 0.45 |
|  | cuddliness | | -0.24 |
|  | soothability | | 0.27 |
| *Orienting/Regulation* | smiling and laughter | | 0.33 |
|  | activity level | | -0.21 |
|  | distress to limitations | | -0.28 |
|  | low intensity pleasure | | 0.70 |
|  | cuddliness | | -0.56 |
|  | duration of orienting | | 0.43 |
|  | soothability | | 0.43 |

**Supplementary Table 2: Descriptive statistics of the MSEL scores at the first 6 months.**

|  | Mean |  | Median | Std. | (Lower) 5% | (Upper) 95% |
| --- | --- | --- | --- | --- | --- | --- |
| E.L.C. | 103.24 |  | 105 | 13.10 | 79.85 | 122.6 |
| E.L. | 55.34 |  | 55 | 8.25 | 42 | 68.15 |
| F.M. | 46.82 |  | 46 | 7.88 | 39.1 | 59.15 |
| G.M. | 49.42 |  | 49 | 8.2 | 36.7 | 61.6 |
| R.L. | 52.45 |  | 51 | 11.67 | 36.85 | 72.6 |
| Vis. | 51.66 |  | 53 | 10.02 | 33.85 | 64.75 |

**Supplementary Table 3: Descriptive statistics of the original 14 IBQ-R scores and the 3 IBQ-R latent factors at the first 6 months.**

|  | Mean | Median | | Std. | | (Lower) 5% | | (Upper) 95% | |
| --- | --- | --- | --- | --- | --- | --- | --- | --- | --- |
| Activity level | 4.3 | 4.47 | | 0.76 | | 3.07 | | 5.4 | |
| Distress to limitations | 3.56 | 3.56 | | 0.87 | | 2.38 | | 4.88 | |
| Fear | 2.22 | 1.87 | | 0.88 | | 1.31 | | 3.86 | |
| Duration of orienting | 3.68 | 3.74 | | 0.85 | | 2.25 | | 4.92 | |
| Smiling and laughter | 4.64 | 4.58 | | 1.12 | | 3.11 | | 6.5 | |
| High intensity pleasure | 5.41 | 5.31 | | 0.85 | | 3.92 | | 6.75 | |
| Low intensity pleasure | 5.31 | 5.38 | | 0.87 | | 4.24 | | 6.45 | |
| Soothability | 4.9 | 4.88 | | 0.73 | | 3.81 | | 5.99 | |
| Falling reactivity | 4.87 | 4.96 | | 0.86 | | 3.54 | | 6.18 | |
| Cuddliness | 6 | 6 | | 0.57 | | 5.13 | | 6.81 | |
| Perceptual sensitivity | 3.45 | 3.41 | | 1.12 | | 2 | | 5.49 | |
| Sadness | 3.5 | 3.43 | | 0.95 | | 2 | | 5.48 | |
| Approach | 4.14 | 4.08 | | 1.08 | | 2.33 | | 5.82 | |
| Vocal reactivity | 4.24 | 4.21 | | 0.92 | | 3 | | 5.72 | |
| SUR | -0.2 |  | -0.14 | | 1.05 | | -1.97 | | 1.63 |
| NEG | 0.13 |  | 0.29 | | 1.04 | | -1.44 | | 1.44 |
| REG | 0.06 |  | 0.06 | | 0.89 | | -1.18 | | 1.61 |

**Supplementary Table 4: Correlation among the 14 nutrients.**

|  | **TSFA** | **TMUFA^2^** | **TPUFA** | **n-6/n-3** | **ARA** | **DHA** | **ARA/DHA** | **PC** | **PE** | **PI** | **SPH** | **Choline** | **Pcho** | **GPC** |
| --- | --- | --- | --- | --- | --- | --- | --- | --- | --- | --- | --- | --- | --- | --- |
| **TSFA** | NA^1^ | ***0.86*** | ***0.69*** | 0.04 | ***0.74*** | 0.35 | -0.20 | 0.27 | 0.18 | 0.13 | 0.21 | -0.23 | 0.22 | -0.04 |
| **TMUFA** | NA | NA | ***0.84*** | -0.01 | ***0.83*** | 0.44 | -0.22 | 0.23 | 0.18 | 0.04 | 0.17 | -0.28 | 0.25 | -0.04 |
| **TPUFA** | NA | NA | NA | -0.09 | ***0.76*** | 0.32 | -0.14 | 0.17 | 0.24 | 0.16 | 0.20 | -0.23 | 0.27 | -0.04 |
| **n-6/n-3** | NA | NA | NA | NA | 0.05 | ***-0.53*** | ***0.51*** | -0.11 | -0.08 | 0.01 | 0.05 | -0.06 | 0.02 | 0.02 |
| **ARA** | NA | NA | NA | NA | NA | 0.34 | -0.04 | 0.23 | 0.19 | 0.07 | 0.18 | -0.23 | 0.31 | -0.02 |
| **DHA** | NA | NA | NA | NA | NA | NA | ***-0.81*** | 0.24 | 0.09 | -0.05 | -0.10 | -0.16 | 0.10 | -0.15 |
| **ARA/DHA** | NA | NA | NA | NA | NA | NA | NA | -0.10 | 0.08 | 0.00 | 0.18 | 0.05 | 0.10 | 0.08 |
| **PC** | NA | NA | NA | NA | NA | NA | NA | NA | ***0.74*** | ***0.63*** | ***0.66*** | -0.27 | 0.08 | -0.18 |
| **PE** | NA | NA | NA | NA | NA | NA | NA | NA | NA | ***0.61*** | ***0.77*** | 0.00 | 0.00 | -0.16 |
| **PI** | NA | NA | NA | NA | NA | NA | NA | NA | NA | NA | ***0.69*** | -0.06 | -0.06 | -0.14 |
| **SPH** | NA | NA | NA | NA | NA | NA | NA | NA | NA | NA | NA | 0.02 | -0.05 | -0.13 |
| **Choline** | NA | NA | NA | NA | NA | NA | NA | NA | NA | NA | NA | NA | ***-0.50*** | 0.08 |
| **Pcho** | NA | NA | NA | NA | NA | NA | NA | NA | NA | NA | NA | NA | NA | 0.08 |
| **GPC** | NA | NA | NA | NA | NA | NA | NA | NA | NA | NA | NA | NA | NA | NA |

^1^ NA: omitted due to matrix symmetry.

^2^ Numbers in bold italic are correlations with p < 0.001.

**Supplementary Table 5: Multiple regression results for MSEL scores/IBQ-R factors with the selected nutrients.**

|  | E.L.C. | G.M. | R.L. | SUR | REG |
| --- | --- | --- | --- | --- | --- |
| R-squared | 0.31 | 0.42 | 0.36 | 0.43 | 0.23 |
| Adjusted R-squared | 0.12 | 0.26 | 0.18 | 0.31 | 0.12 |
| Raw p-value^1^ | 0.05 | 0.05 | 0.005 | 0.0008 | 0.017 |
| Adjusted p-value^2,3^ | 0.05 | 0.11 | **0.025** | **0.003** | **0.03** |
| TSFA^4,5,6^ | -0.42 (0.03) | - | -0.30 (0.05) | - | - |
| TPUFA | - | - | - | - | - |
| TMUFA | - | -0.51 | - | - | - |
| ARA | - | - | - | 0.42 (0.008) | 0.39 (0.02) |
| n-6/n-3 | 0.26 (0.19) | 0.19 | 0.37 (0.006) | - | - |
| DHA | 0.55 (0.02) | 0.30 | 0.75 (0.002) | - | - |
| ARA/DHA | - | - | - | - | - |
| PC | - | - | - | - | - |
| PE | - | - | 0.30 (0.07) | - | - |
| PI | - | - | - | 0.19 (0.24) | - |
| SPH | -0.22 (0.23) | -0.24 | - | - | - |
| Choline | - | - | - | - | - |
| PCho | - | - | - | 0.27 (0.08) | - |
| GPC | - | - | - | - | - |
| Age | -0.006 (0.98) | 0.28 (0.12) | 0.30 (0.03) | 0.18 (0.26) | 0.18 (0.31) |
| Sex (if Male) | 0.03 (0.87) | 0.08 (0.61) | -0.01 (0.79) | -0.03 (0.83) | -0.004 (0.98) |
| Site (if UNC) | 0.20 (0.29) | 0.44 (0.01) | 0.10 (0.63) | -0.32 (0.05) | 0.22 (0.21) |
| Income (if < 75k) | -0.13 (0.44) | 0.14 (0.35) | -0.15 (0.21) | -0.15 (0.34) | 0.06 (0.71) |

^1^ ANOVA F-test comparing to the reduced model (confounders only); results with raw p > 0.05 are not shown.

^2^ P-values in this line indicate the p-values after FDR correction for multiple comparison.

^3^ Adjusted p-values less than 0.05 are shown in bold text.

^4^ Numbers in parentheses show the p-values of the coefficients using the t-test statistic.

^5^ Hyphen “-” indicates that this variable is not selected in this model.

^6^ Coefficients here are standardized linear regression coefficients.

#### 2.2 Supplementary Figures

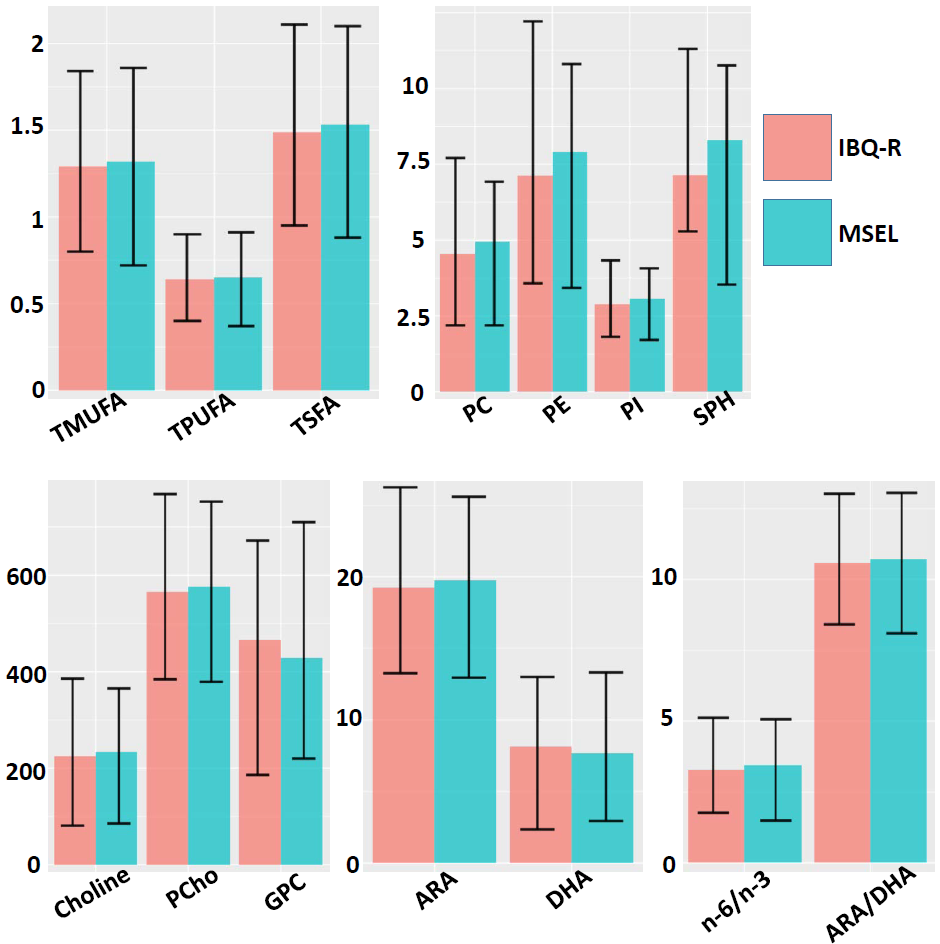

**Supplementary Figure 1.** Barplots of the HM nutrients (Mean ± STD) among the 38 subjects (blue) with MSEL and 42 subjects (red) with IBQ-R measurements at the first 6 months.

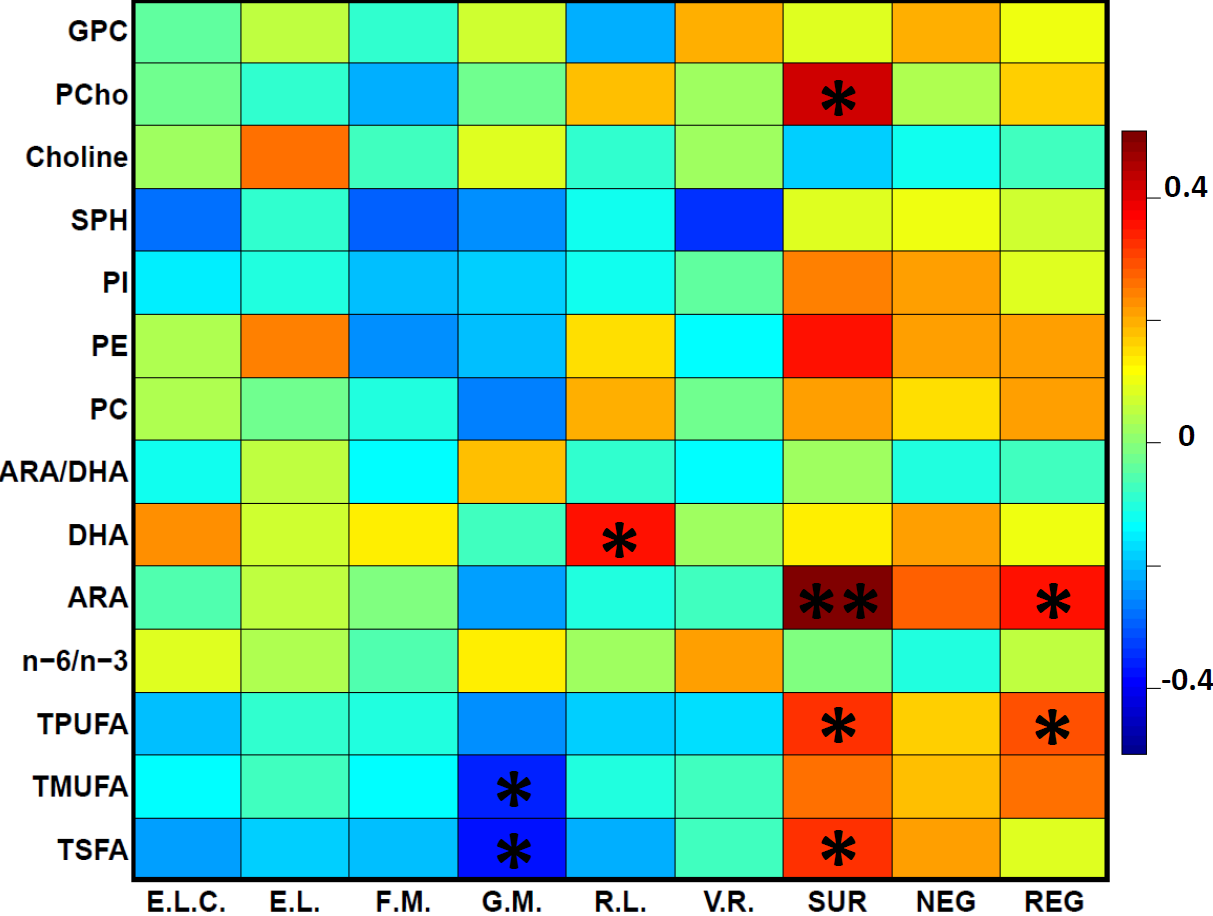

**Supplementary Figure 2**. The heatmap of the results shown in Table 3 in the main text, which shows the marginal associations of each nutrient with each subdomain of MSEL and IBQ-R. Confounding factors age, sex, site and household income (if < 75k) were controlled. The asterisks indicate raw p < 0.05 and double asterisks indicate adjusted p < 0.05.

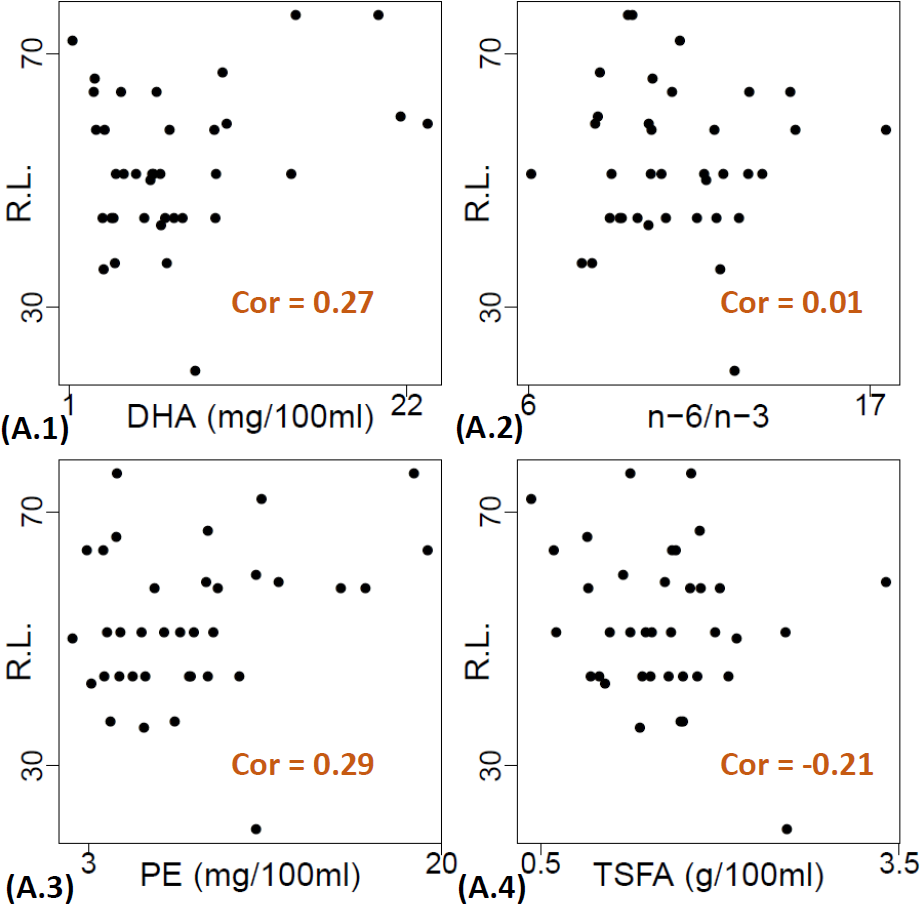

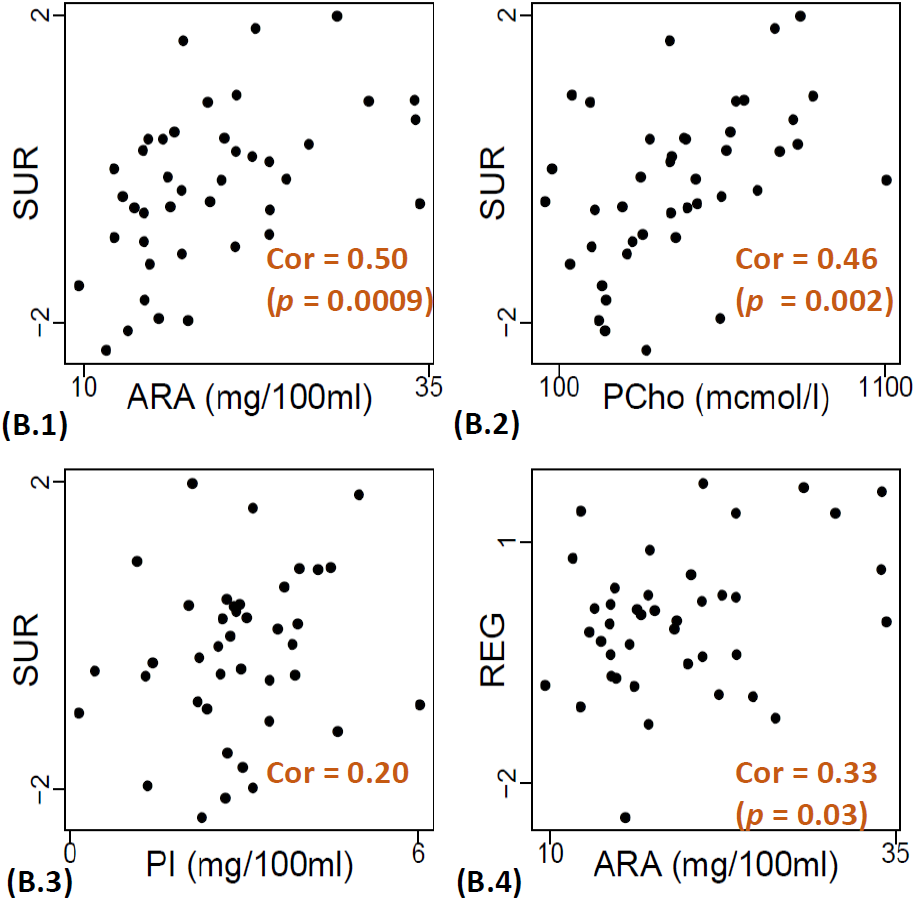

**Supplementary Figure 3.** The scatterplots of selected nutrients for receptive language score (R.L.), surgency (SUR) and regulation (REG) are shown in (A.1)-(A.4), (B.1)-(B.3), and (B.4) respectively.
